# HD2Net: A Deep Learning Framework for Simultaneous Denoising and Deaberration in Fluorescence Microscopy

**DOI:** 10.1101/2025.01.06.631475

**Authors:** Xuekai Hou, Yue Li, Chad M. Hobson, Hari Shroff, Min Guo, Huafeng Liu

## Abstract

Fluorescence microscopy is essential for biological research, offering high-contrast imaging of microscopic structures. However, the quality of these images is often compromised by optical aberrations and noise, particularly in low signal-to-noise ratio (SNR) conditions. While adaptive optics (AO) can correct aberrations, it requires costly hardware and slows down imaging; whereas current denoising approaches boost the SNR but leave out the aberration compensation. To address these limitations, we introduce HD2Net, a deep learning framework that enhances image quality by simultaneously denoising and suppressing the effect of aberrations without the need for additional hardware. Building on our previous work, HD2Net incorporates noise estimation and aberration removal modules, effectively restoring images degraded by noise and aberrations. Through comprehensive evaluation of synthetic phantoms and biological data, we demonstrate that HD2Net outperforms existing methods, significantly improving image resolution and contrast. This framework offers a promising solution for enhancing biological imaging, particularly in challenging aberrating and low-light conditions.

## 1. Introduction

Fluorescence microscopy is an important tool for biological research, enabling imaging of biological phenomena with molecular contrast [1]. However, imaging quality can be compromised by several factors [2], including optical aberrations. Aberrations distort the wavefront of light, leading to suboptimal focusing and reduced image resolution, contrast, and signal [3],[4]. This effect becomes more pronounced as imaging depth increases, complicating the visualization of structures within thick biological samples.

Adaptive optics [5],[6](AO) mitigates aberrations by measuring the distorted wavefront and subsequently canceling it by applying a corrective wavefront, thereby restoring image quality. However, AO system require specialized and expensive components, such as deformable mirrors and wavefront sensors, making them challenging to incorporate and maintain. Additionally, the wavefront measurement process slows down imaging and introduces more illumination dose [7], limiting the practicality of AO in dynamic biological studies.

To address these challenges, we previously introduced the DeAbe [8] method, which employs deep learning techniques to ameliorate image degradation caused by aberrations. DeAbe introduces synthetic aberrations to images acquired on the shallow side of image stacks so that the synthetically degraded images mimic those acquired deeper into the volume; trains a neural network to reverse the effect of these aberrations given that the original shallow side images are near-diffraction-limited; and then applies the trained neural network to data acquired deeper into the volume. Through this carefully designed computational strategy, DeAbe eliminates the need for complex wavefront measurements, significantly reducing costs and simplifying image acquisition. DeAbe offers substantial enhancements in image quality and subsequent downstream analyses when applied to images of thick samples, such as improved assessment of vascular directionality in mouse tissues and better membrane/nuclear segmentation in *C. elegans* embryos. DeAbe is also applicable across a range of microscopy modalities, including confocal, light sheet, multiphoton, and super-resolution microscopy.

Despite its advantages, DeAbe faces limitations when dealing with low signal-to-noise ratio (SNR) conditions, where the impact of noise on image quality becomes increasingly detrimental. In fluorescence microscopy, the “pyramid of frustration” dilemma arises, where researchers must balance phototoxicity/photobleaching, observation duration, and resolution[9]. These tradeoffs are particularly salient when considering live imaging, when reducing the illumination light dose may be essential, yet the resulting loss in SNR [10] may hinder image restoration. Some efforts [11] have drawn attention to the noise influence but are limited to 2D models and lack a thorough exploration of the noise effects.

To enhance the performance of DeAbe in low SNR scenarios, we propose a Hybrid Denoising and Deaberration Network (HD2Net). This self-supervised deep learning approach integrates noise estimation into the existing DeAbe framework, enabling simultaneous denoising and aberration correction. Here we show this advancement improves image restoration in challenging, low-light imaging conditions. We demonstrate the effectiveness of HD2Net in enhancing the quality of fluorescence microscopy on diverse samples, including phantom objects, fixed cells, and *C. elegans* embryos labeled with membrane and nuclear markers.

## 2. Methods

### 2.1 Overall framework of HD2Net

Our method is based on two key observations: 1) Images located on ‘shallow’ or ‘near side’ of the 3D fluorescence volumes are typically near-diffraction limited without pronounced aberrations [12]. 2) As imaging depth increases, signal degrades due to light distortion and absorption, making the effect of aberrations and noise in the deeper regions more pronounced compared to the superficial sections [13]. Based on these two insights, we use the high-quality ‘shallow’ images as a ground truth reference for generating data that mimics the degradation observed in images located in the deeper part of the sample, enabling us to train HD2Net for de-aberration. To achieve simultaneous denoising and aberration suppression, the overall workflow is shown in **Fig 1**. This includes synthetic data generation, network training, and network inference.

**Fig. 1.**
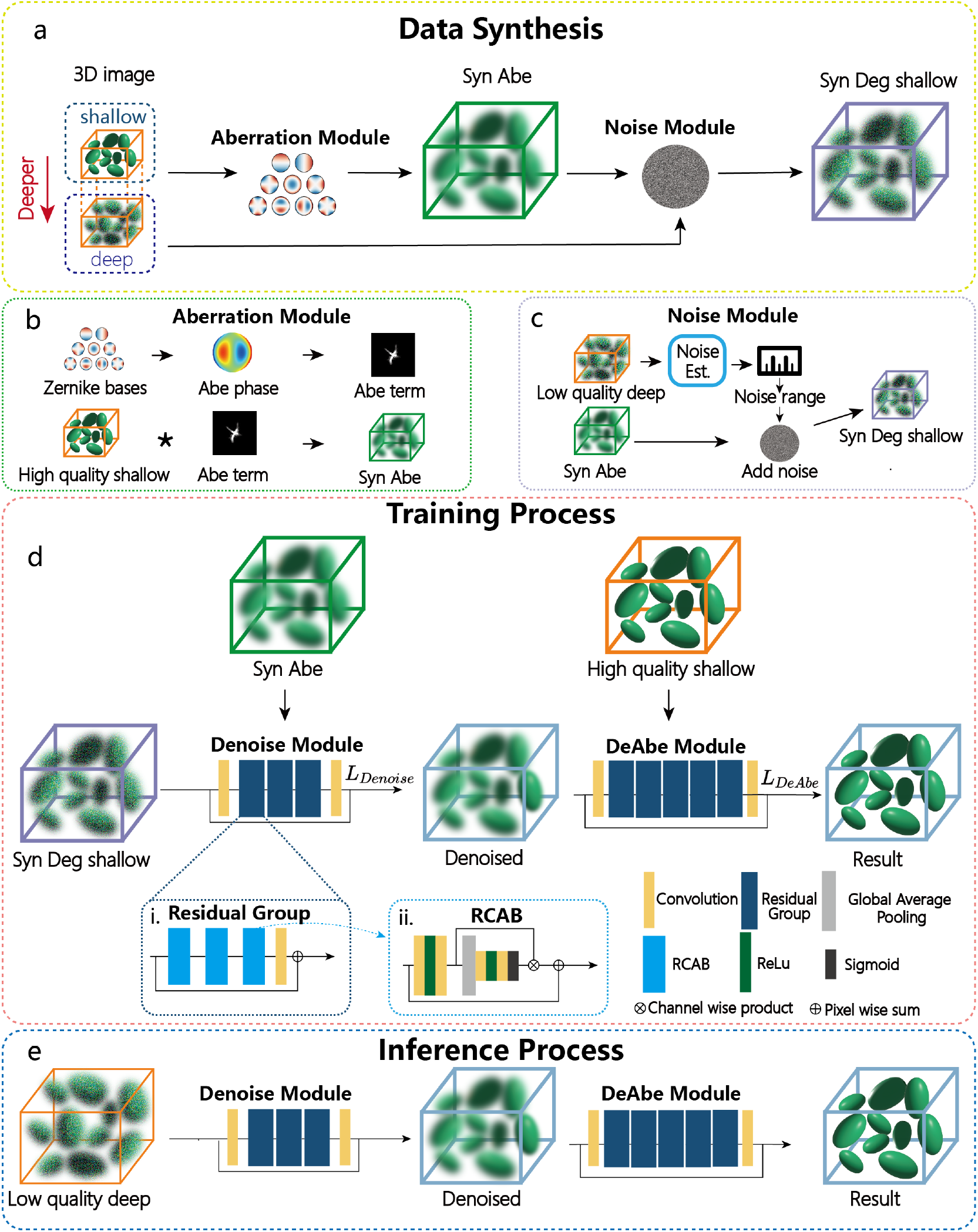
Overall framework of HD2Net. **a)** Data synthesis pipeline. High-quality ‘shallow’ subvolumes are extracted from the acquired 3D volume data and passed through the Aberration and Noise modules to obtain the synthetic degraded data (Syn Deg shallow). The degradation is designed so that the synthetic data resembles the low-quality ‘deep’ regions in the acquired volumes. **b)** The Aberration Module: Zernike basis functions (Zernike bases) and their associated coefficients are used to obtain the aberrated wavefronts (Abe phase). The aberration term (Abe term) is derived from the aberrated phase through image formation process (see **Methods 2.3**). The ‘shallow’ images are convolved with the aberration term to obtain the synthetic aberrated images (Syn Abe). **c)** The noise estimation model (Noise Est.) estimates the noise level from the low quality ‘deep’ images. Noise is added to the Syn Abe images based on the estimated noise level to produce the degraded synthetic (Syn Deg shallow) images. See **Methods** for more details. **d)** Training in HD2Net consists of successively training Denoise and DeAbe Modules. The degraded images (Syn Deg Shallow) are used in conjunction with Syn Abe data to train the Denoise module. The resulting denoised images (Denoised) are then used in conjunction with High quality shallow data to train the DeAbe module. The structures of the modules are illustrated using boxes with different colors, **i** and **ii** show the detailed structures of the Residual Group and RCAB (residual channel attention blocks), respectively. **e)** Inference process using HD2Net. Once the network is trained, experimentally acquired, low quality deep volumes can be fed into the network to suppress the effect of aberrations and noise.

### 2.2 General training data synthesis

We first extracted the shallow and deep subvolumes from experimentally acquired volumetric data **(Fig. 1a)**. The shallow subvolume is defined by the planes nearest to the detection objective which have no or minor aberrations and that possesses high SNR, as described in our previous paper [8]. For simulated phantoms, the shallow subvolumes are defined by synthetic aberration-free volumes. Similarly, the deeper subvolumes consist of image stacks with obviously degraded signal and resolution. To better simulate the degradation process, we use Aberration and Noise Modules to incrementally introduce aberrations and noise to the shallow subvolumes. The degradation is inspired by the incoherent image formation process:

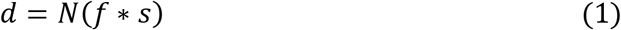

where *f* is the object; *d* is the observed, degraded data; *s* is the incoherent point spread function (PSF); * is the convolution operation; and *N*(⋅) applies Poisson noise, which is usually dominant in fluorescence microscopy.

### 2.3 Aberration synthesis

We use the Aberration Module to generate the noise-free synthetic aberrated images (Syn Abe). Aberration is introduced through the convolution operation, and the specific process is shown in **Fig. 1b**.

The convolution operation in the spatial domain can be transformed into product operation in frequency domain. Thus, the imaging formation formula in the frequency domain can be written with the Fourier transform (ℱ(⋅)) as:

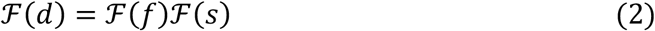

Here, ℱ(⋅) is the Fourier transform, ℱ(*s*) is known as the optical transfer function (OTF). When imaging without aberration, we have:

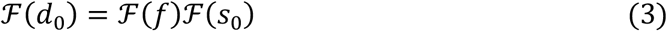

Here *s*_0_ is the ideal PSF. To obtain the aberrated data *d* from the aberration-free data *d*_0_ and eliminate the object term, we substitute Eq.(2) into Eq.(3) to obtain the aberration term τ:

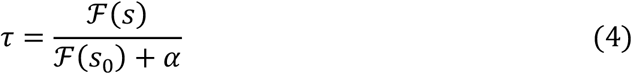

where α is a small value to prevent division by zero. Empirically, it is good when α ≤ 10^−2^, and for the datasets in this paper, α is set to 10^−6^. We can then obtain the aberrated data *d* as:

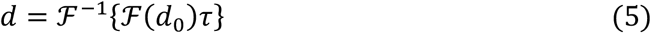

The level and type of aberration is determined by the incoherent PSF *s*. It is given by:

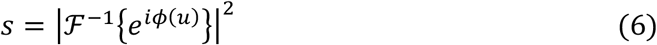

which is applicable to signals confined within the pupil aperture. The aberrated phase term *ϕ*(*u*) can be expressed using the combination of a series of Zernike basis functions *ϕ*_*m*_(*u*) and associated coefficients *c*_*m*_:

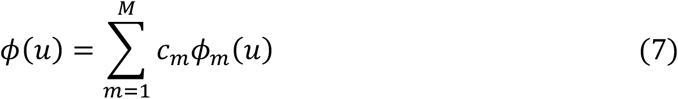

*ϕ*_*m*_(*u*) describes the different aberration modes, *c*_*m*_ indicates the magnitude of the aberrations. The degree of aberration can thus be adjusted by modifying *ϕ*_*m*_(*u*) and *c*_*m*_. We used the ANSI [14] convention when indexing the Zernike coefficients and added aberrations up to the 4^th^ Zernike order (*m*_*max*_= 14), except for piston and tilt components (*m* = 0, 1, 2).

### 2.4 Noise synthesis

The noise synthesis workflow is shown in **Fig. 1c**. To determine the amount of noise that needs to be added, we use a Noise Module to estimate the noise level in the ‘deep’ low-quality subvolumes. Under low-light conditions, the noise distribution in fluorescence microscopy imaging follows a Poisson distribution [15].

Using variance-stabilizing transformations[16],[17] (VST) can convert signal-dependent Poisson noise to additive Gaussian noise, removing the data dependency of the noise. One of the most popular variance-stabilizing transformations is the Anscombe transform [18].

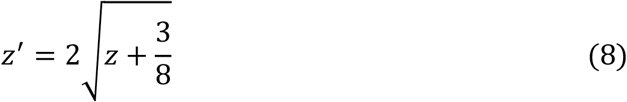

where *z* represents the observed images, and *z*′ the transformed images. After the Anscombe transform, we can write the imaging model as:

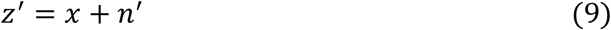

where *x* is the noise-free image, *n*′ is the transformed Gaussian noise. There are many methods for Gaussian noise estimation. Here, we use Liu’s method[19] which can estimate the noise level from image patches using weak textured area selection. Using Principal Component Analysis (PCA) [20] on Eq. (9), we can derive the following equation:

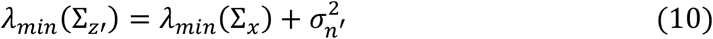

where Σ_*z*′_ is the covariance matrix of the noisy image z’, Σ_*x*_ denotes the covariance matrix of the clean image x, λ_*min*_ (Σ) represents the minimum eigenvalue of the matrix Σ, and σ_n′_ is the standard deviation of the transformed Gaussian noise and represents the noise level. It is difficult to estimate the unknown *σ*_*n*′_, because the minimum eigenvalue of the covariance matrix from noise-free patches is also unknown. Liu et al [19] addressed this issue by an iterative framework to select weak textured patches form the noisy image. The minimum eigenvalue of the covariance matrix of these patches is approximately zero. Therefore, the noise level can be estimated simply as:

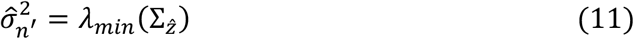

where 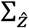 is the covariance matrix of the selected weak textured patches. After obtaining the noise level parameter *σ*_*n*′_, we use the following method to add Poisson noise to the noise-free aberrated images:

Because the Poisson noise arises due to intrinsic noise from the signal, we first enhance or weaken the signal intensity of the images obtained from the Aberration Module, resulting in a series of images with different signal intensities. Then we use MATLAB’s *addnoise* function to introduce Poisson noise to these images and obtain a series of images with varying noise levels. We use the noise estimation module to estimate the noise level for these images by applying Equations (8-11) as described above. Finally, we retain the data (Syn Deg shallow) that matches the target noise level *σ*_*n*′_.

### 2.5 HDN2Net construction and implementation

The HDB2Net network (**Fig. 1d**) consists of a Denoise Module and a DeAbe Module. Denoising and deaberration are executed sequentially, in a process that can be described as:

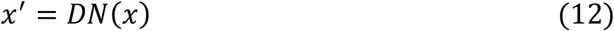

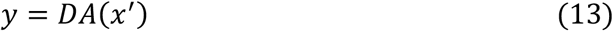

where *x* is the synthetic degraded raw image, *x*′ is the denoised result which contains aberrations, *y* is the deaberrated result. *DN* represents the Denoising Module and *DA* represents the DeAbe Module.

We adapted the three-dimensional residual channel attention networks (3D-RCAN)[21] (https://github.com/AiviaCommunity/3D-RCAN) to construct both the Denoising Module and the DeAbe Module. The basic structure of 3D-RCAN [22] is the Residual Group (RG), with each RG containing 3 residual channel attention blocks (RCAB). RG can be described as:

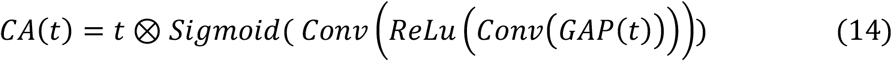

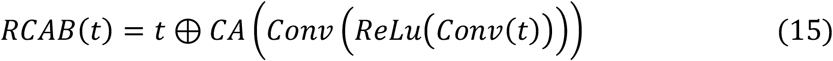

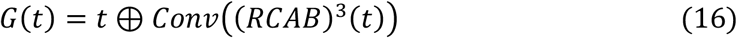

where *CA* is the channel attention module, *CONV* represents the convolution layer, *GAP* is the global average pooling, *ReLu* and *Sigmoid* are the activation functions,⊗ is the channel-wise multiplication, ⊕ is the pixel-wise summation. Since deaberration is more complex than denoising, we added more RGs for the DeAbe module [21], using 3 RGs for the Denoising Module and 5 RGs for the DeAbe Module:

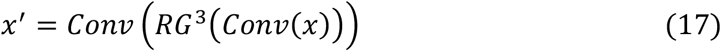

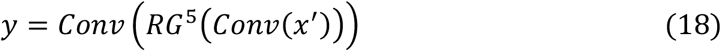

The number of Convolution layers for feature extraction is 32; the reduction ratio for CA is set to 8. During the network training process, we used the Syn Abe data to supervise the Denoising Module and the high quality shallow images to supervise the whole network. This separated supervision approach aims to enhance the denoising and deblurring performance of the network by incorporating more image details than single-step supervision by only the high quality shallow images (**Fig. 1d**). We used the Charbonnier loss [23] in training both modules:

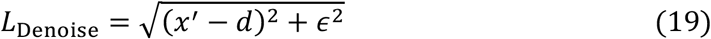

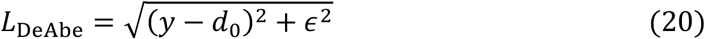

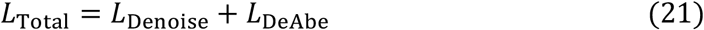

where *d* is the Syn Abe images, *d*_0_ is the high-SNR aberration-free images. ϵ is a small constant and set to 10^−3^ which is same as Charbonnier’s setting [23].

During training, the number of iterations is set to 200000, the training patch size is set to 64 × 64 × 64, the learning rate is set to 10^−4^. These parameters are determined based on our empirical experience. Generally, the number of iterations should not be too small to prevent the network from being underfitted. When applying the model, we also set patch size as 64 × 64 × 64. For image volumes larger than this patch size, we split them into patches, applied the network to each patch, and then stitched the patches back using linear blending to minimize boundary artifacts. The training and model application were performed within Python 3.9.18 on a Linux workstation with the following specifications: CPU - Intel(R) Xeon(R) CPU E5-2680 v3 @ 2.50GHz, RAM - 128 GB, GPU - NVIDIA GeForce TITAN RTX with 24 GB memory.

For quantitative analysis, we used structural similarity index (SSIM) and peak signal-to-noise ratio (PSNR) referenced to the ground truth images to evaluate the network outputs. The PSNR was computed as:

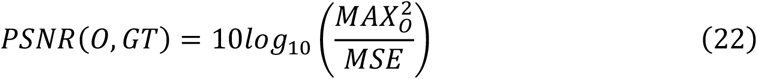

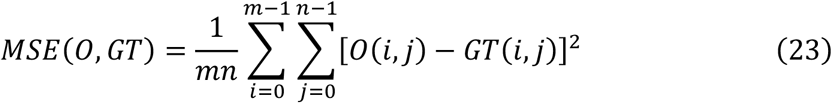

where *O* is network outputs and *GT*is ground truth. *MAX*_*O*_ is the maximum possible pixel value of the image. When the pixels are represented as normalized pixels, this is 1. A higher *PSNR* value means the network output is more similar to the ground truth.

The SSIM was computed as:

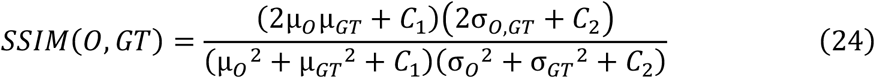

where *µ*_*GT*_, *µ*_*O*_ are the mean values of the *GT* and *O*; *σ*_*GT*_^2^, *σ*_*O*_^2^ are the variances of the *GT* and *O*; *σ*_*O,GT*_is the covariance of *GT* and *O*; and *C*_1_ and *C*_2_ are small constants that prevent the denominator from becoming zero (here *C*_1_ = 1*e*^−4^ and *C*_2_ = 9*e*^−4^). A higher *SSIM* value means the network output is more similar to the ground truth.

## 3. Results

### 3.1 Performance on simulation data

We conducted a series of simulation experiments to validate the advantages of the HD2Net pipeline over the original DeAbe pipeline[8]. We generated three-dimensional phantom volumes with points, lines, spheres, circles, and spherical shells in random directions and positions as the ground truth (GT)[24]. Then we degraded them to generate the inputs for training and testing. Compared with the original data synthesis in DeAbe, our new pipeline further considered the impact of noise. To verify the function of the Noise Module, we adopted two degradation strategies, one with the Noise Module (HD2Net) and the other without the Noise Module (HD2Net, w/o NM). The training inputs were mixtures of different aberration levels and noise levels. Since Poisson noise is expected to be dominant in fluorescence microscopy, we define the SNR as 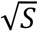,where *S* is the signal defined by the average of all pixels with intensity above a threshold (here set as 1% of the maximum intensity of the blurred objects in the noise-free image). The aberrations ranged from 1 to 4 radians. The SNR is set to 20.7 without the Noise Module, and between 1.9 and 20.7 with the Noise Module. During testing, the trained models were applied to specific noise levels.

To demonstrate the improvement of HD2Net over DeAbe, we used the data synthesis strategy to generate phantom training datasets by degrading shallow subvolumes with aberrations only (i.e. training datasets without using the Noise Module) and trained each network. Then we obtained DeAbe model and HD2Net (w/o NM) model respectively, and compared their performance by applying each model on the same test data containing a significant amount of noise (**Fig. 2a**, SNR=1.9). Due to the misalignment between the SNR in training vs. test data, both DeAbe and HD2Net (w/o NM) failed to achieve optimal restoration. Nevertheless, the HD2Net (w/o NM) results contain less noise and structural deformation than the DeAbe result, demonstrating that the Denoising Module in HD2Net architecture can suppress noise amplification to some extent. We further considered the noise synthesis during data generation and used data with degraded by noise and aberrations to train the HD2Net (i.e., training a full denoising and deaberration model). In this case, HD2Net performed well both laterally and axially, with the shapes of circles and lines being well preserved and closely resembling the Ground Truth (GT). This demonstrates the value of explicitly accounting for the impact of noise during data generation.

**Fig. 2.**
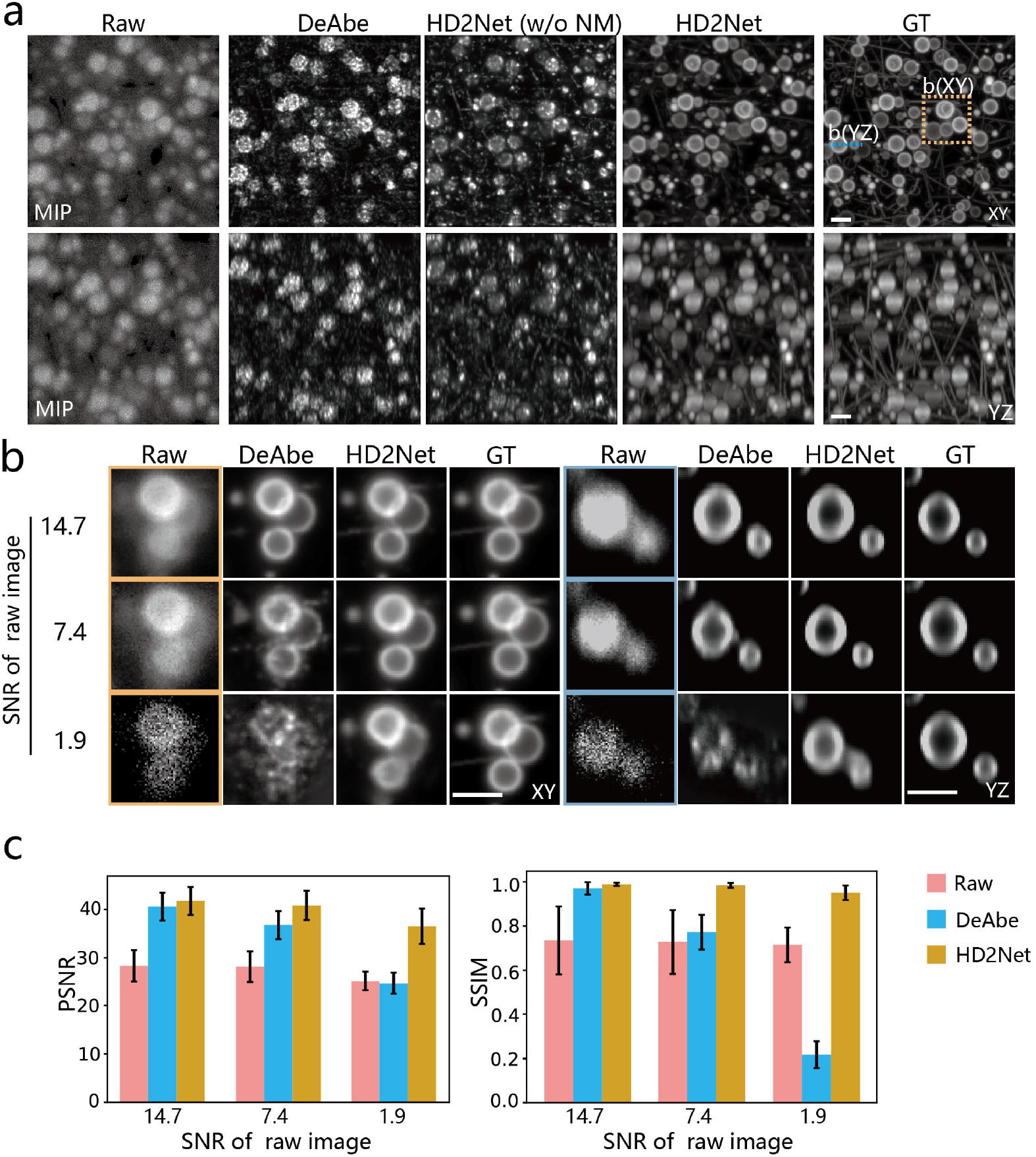
HD2Net outperforms DeAbe on synthetic data with different SNR. **a)** Lateral and axial maximum intensity projection (MIP) of images degraded with large amounts of noise and aberrations (SNR=1.9), comparing low-quality raw image, the restoration provided by using DeAbe, the restoration result of HD2Net trained with noise-free data (HD2Net w/o NM), the restoration result of HD2Net trained by data with Noise Module (HD2Net), and the ground truth (GT). The first row shows the lateral results, and the second row shows the axial results. **b)** The restoration quality as a function of input data with different SNR, comparing raw image, the restoration result of DeAbe, the restoration result of HD2Net, and GT. The left four columns show the lateral views of yellow rectangle in **a**. The right four columns show the axial views taken across the cyan dashed line in **a** Rows from top to bottom correspond to low noise (SNR = 14.7), medium noise (SNR = 7.4), and high noise (SNR = 1.9) conditions respectively. **c)** Quantitative analysis with PSNR and SSIM for **a** and **b**, indicating the good performance of HD2Net. Means and standard deviations are obtained from N = 50 simulations (10 independent phantom volumes, each aberrated with 5 randomly chosen aberrations). Scalebars: 3.5 µm in **a** and **b**.

Next, we compared H2DNet to DeAbe under different SNR conditions. DeAbe was trained with data generated without the Noise Module and HD2Net was trained with data generated with the Noise Module. The test data included low, medium, and high levels of noise, with SNR being 14.7, 7.4 and 1.9 respectively. When the SNR of the test data was high, both DeAbe and HD2Net performed well. However, the performance of DeAbe decreased rapidly as noise increased (**Fig. 2b**), whereas the performance of HD2Net only decreased slightly, and remained similar to the GT results. These qualitative visual impressions were also consistent with quantitative analysis via SSIM and PSNR metrics against the GT reference (**Fig. 2c**). In all cases, HD2Net outperformed DeAbe, as assessed via quantitative metrics. Although the PSNR and SSIM of HD2Net and DeAbe both decline with increased noise, DeAbe’s PSNR drops from 40.9dB to 25.7dB, while HD2Net’s PSNR remains above 36.8dB. DeAbe’s SSIM decreases from 0.97 to 0.22, whereas HD2Net’s SSIM stays above 0.94. These results confirm that including the Noise Module during training data synthesis helps the proposed workflow better cope with noise, achieving superior performance.

### 3.2 The impact of different noise addition strategies

Using the Noise Module in training data synthesis is a key factor in improving performance over DeAbe. We also used simulations to help inform how much noise should be added to the training dataset. Since the SNR is affected by both aberrations and noise, we used the factor noise level (NL) defined previously [19] for our assessment to isolate the effect of noise. The noise level represents the standard deviation of noise intensity in the normalized image, with higher values indicating greater noise and lower SNR. In these experiments, we added random aberrations with a fixed magnitude of 2 radians and adjusted the noise conditions by adjusting the parameters of the Noise Module.

Since the training data were synthesized with a range of noise levels, we first studied the impact of the noise range (NR) in training data on the network performance in test data with fixed noise levels (**Fig. 3a**). We set three different noise levels for the test data (Test NL): low (5.5 × 10^−3^, equivalent to SNR of 20), medium (11.1 × 10^−3^, equivalent to SNR of 10) and high (26.3 × 10^−3^, equivalent to SNR of 2). For each noise level in the test data, we set three noise ranges (NR1 (90%-110% of Test NL), NR2 (75%-125% of Test NL), and NR3 (60%-140% of Test NL)) to generate three training datasets, and trained HD2Net with each training dataset to obtain corresponding models. Then each model was applied to test data degraded with noise set at the corresponding noise level. Quantitative analyses through SSIM indicate that the selection of noise ranges under different noise levels does not have a great impact on the results, although using a larger NR shows a slight advantage over using a smaller NR (**Fig. 3a**).

**Fig. 3.**
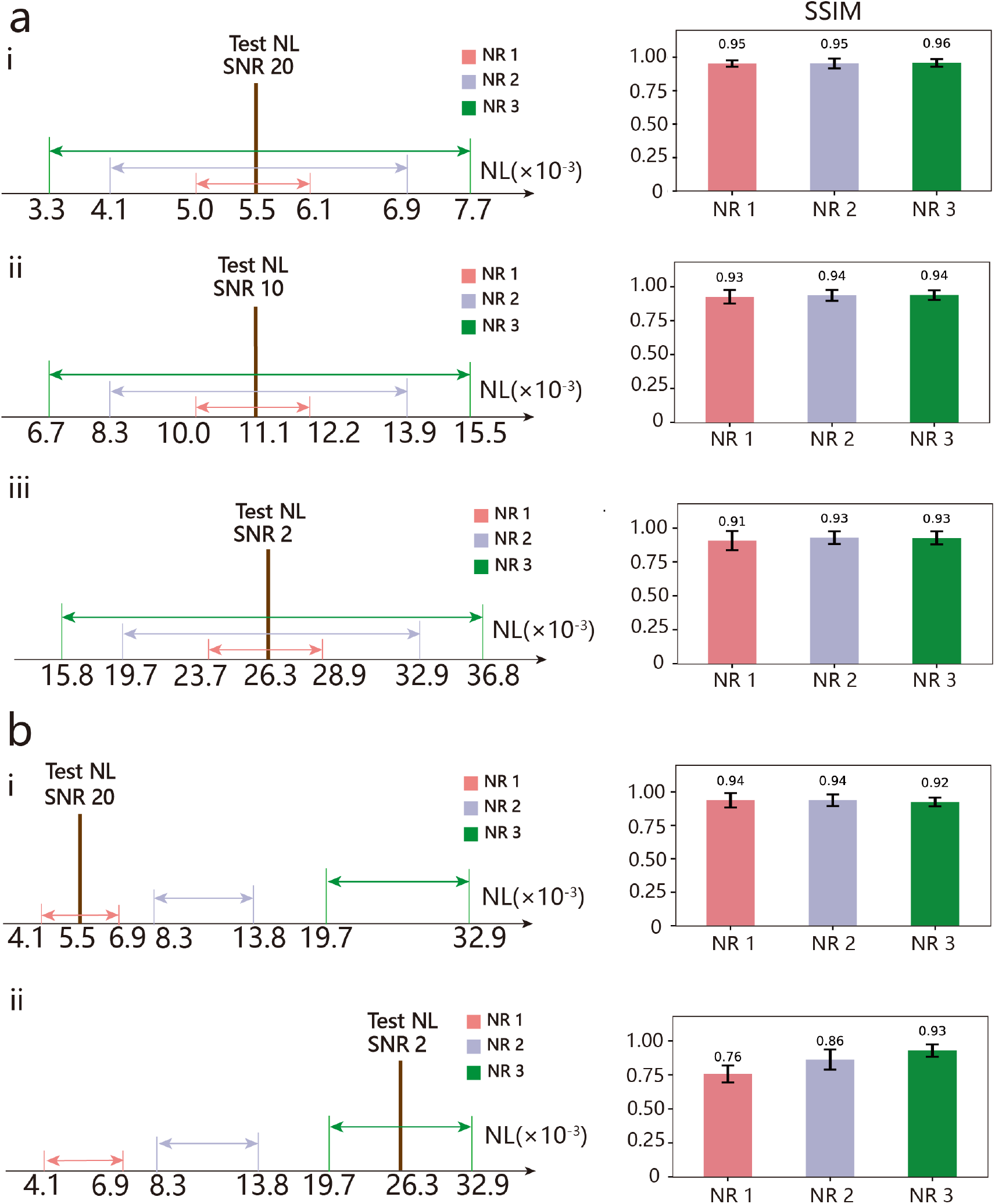
Quantitative analysis of HD2Net under different noise addition strategies. **a)** Influence of noise range (NR) in training data on test data with different noise levels (NL): **(i)** for the low Test NL condition (equivalent to SNR of 20), **(ii)** for the medium Test NL condition (equivalent to SNR of 10), **(iii)** for the high Test NL condition (equivalent to SNR of 2). The vertical lines (brown) perpendicular to the NL axis indicate the noise level of the test data (Test NL). Horizontal lines with different colors indicate the noise range (NR) added to the training dataset, red lines are NR 1 from 90% to 110% of Test NL, gray lines are NR 2 from 75% to 125% of Test NL, green lines are NR from 60% to 140% of Test NL. SSIM analyses are used to evaluate the restoration results. **b)** The impact of the noise level misalignment between the training dataset and the test data: **(i)** for the Test NL (equivalent to SNR of 20) matching training data with a low NR, **(ii)** for the Test NL (equivalent to SNR of 2) matching training data with a high NR. The vertical lines (brown) perpendicular to the NL axis indicate the test NL. Horizontal lines with different colors indicate different NRs of training dataset, red lines are low training NR, gray lines are medium training NR, green lines are high training NR. SSIM analyses are used to evaluate the influence of misalignment. Means and standard deviations from N = 20 measurements are shown.

Next, we investigated the impact of misalignment between the training dataset and the test data on the network’s restoration capability, i.e., the difference in network performance when the noise added to the training set is higher or lower than that of the test set.

In the case of a low noise level in the test data (equivalent to SNR of 20), we set three different training noise levels: training NL matching the testing NL, training NL moderately higher than testing NL, and training NL much higher than testing NL. In the case of high noise levels in the test data (equivalent to SNR of 2), we also set three different training noise levels: training NL matched the testing NL, training NL moderately lower than testing NL, and training NL much lower than testing NL. Unsurprisingly, we found that the restoration will be better when the misalignment between test NL and training NL is small in both cases (**Fig. 3b**). In the case of low NL in the test data, the differences in performance were not obvious; whereas in the case of high NL in the test data, mismatching of the noise levels resulted in a significant performance decrease.

Therefore, when synthesizing training data corresponding to real biological samples, we should ideally adjust the Noise Module according to the noise condition of the deeper subvolumes so that the noise level in training and test data closely match. However, this requires repeated noise estimation during data synthesis. Because the model trained with high NL training data also performs well on low NL test data, in practice we add more noise to the training data and set the noise range to be larger to simplify the data generation process.

### 3.3 Simultaneous denoising and deaberration improves image quality on real biological samples

We next demonstrate the effectiveness of HD2Net on experimentally-acquired image data from biological samples. We first compared HD2Net and DeAbe on PtK2 cell data acquired using lattice light-sheet microscopy equipped with adaptive optics[25],[26] (AO-LLSM). Actin in these cells was immunostained by Phalloidin Alexa Fluor 488. By adjusting the illumination intensity, we can control the noise level in the data. By leveraging the deformable mirror in the AO system, we can introduce aberrations to blur the fine actin mesh at the cell periphery, filamentous actin, and stress fibers. Additionally, we can obtain a low-noise aberration-free experimental reference using the same setup, to benchmark the restoration results. The full details of the sample preparation and imaging experiments can be found in our previous work [8]. The training dataset was derived from the aberration-free image volumes with high illumination intensity via the data synthesis procedure (**Fig. 1**). HD2Net and DeAbe models were trained to restore the images. We find that although HD2Net and DeAbe both improve image resolution and contrast, the noise reduction offered by HD2Net produces a prediction closer to the high SNR, aberration-free reference based on visual inspection (**Fig. 4a**) and line profiles through closely distributed actin fibers (**Fig. 4b**). Additional analysis of the Fourier spectrum shows that the results from HD2Net are very close to those high SNR aberration-free images (**Fig. 4a**, inset).

**Fig. 4.**
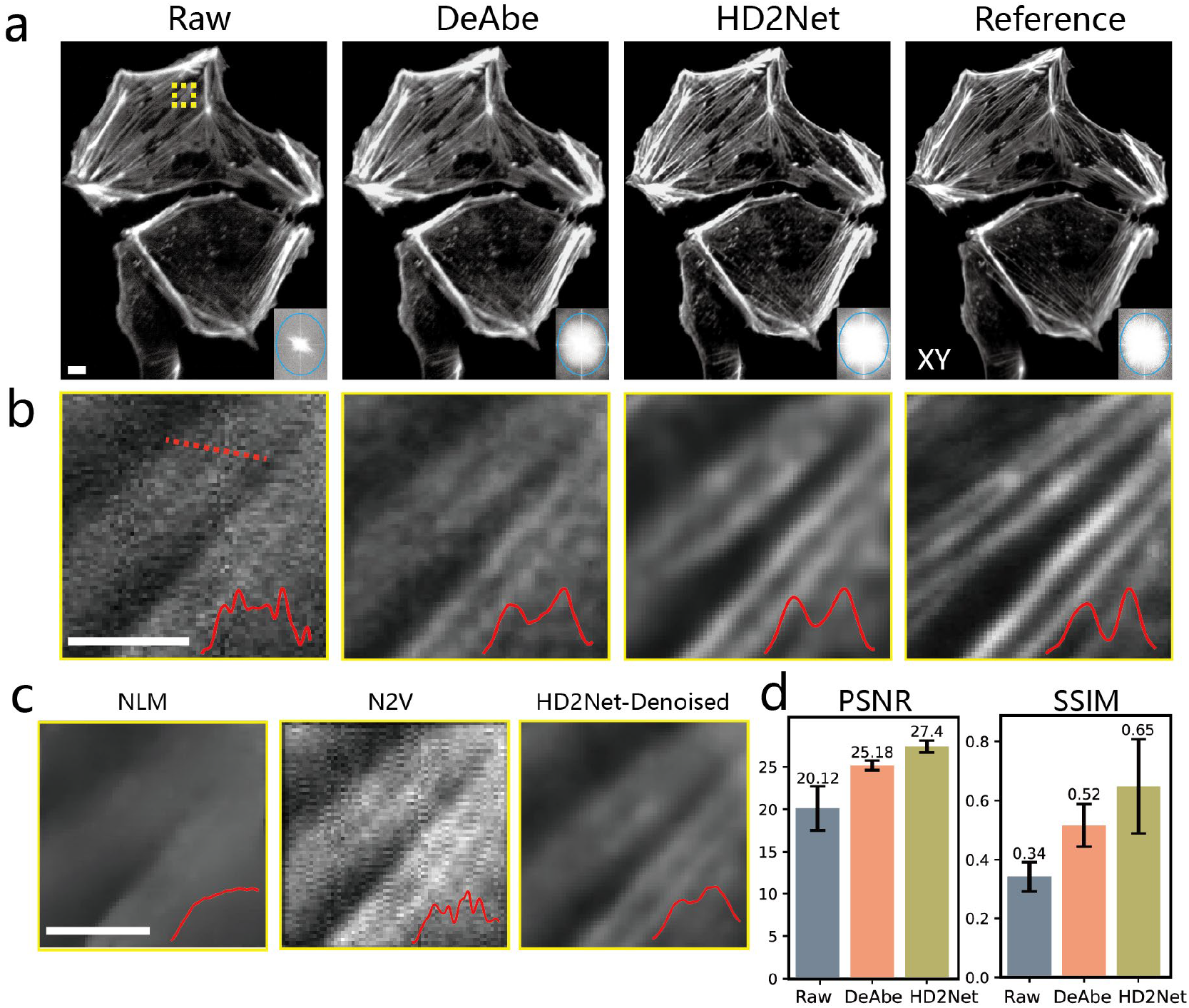
HD2Net outperforms DeAbe and denoised methods on experimentally-acquired data. Fixed Ptk2 cells were stained for actin using Phalloidin Alexa Fluor 488 and imaged with an AO lattice light-sheet microscope. **a)** Left to right: the raw image, the DeAbe model prediction, the HD2Net model prediction, and the low-noise aberration-corrected experimental reference (using 10x more excitation laser power than raw images, aberration-free). The insets on the lower right corner of each image show the Fourier transforms, with the blue ellipse having a horizontal extent of 1/500 nm^−1^ and a vertical extent of 1/400 nm ^−1^. Note that the images have been rotated to be viewed perpendicularly to the coverslip surface, resulting in anisotropic resolution in the lateral plane and the display contrast of raw images has been adjusted to enhance the brightness for visual purpose. **b)** Higher magnification views of yellow rectangular regions in **a**. The insets represent the line profile along the red dashed line. **c)** Denoising the region shown in **b**, comparing Non-Local Means (NLM), Noise2Void (N2V) and HD2Net’s Denoise Module, the insets represent the profile indicated by the same line as in **b. d)** Quantitative comparisons for the restorations shown in **a** using PSNR and SSIM. Means and standard deviations for 12 images are shown. Scalebars: 3.5 µm in **a, b** and **c**.

Next, we compared the Denoise Module in HD2Net with traditional denoising algorithms (Non-local means[27], NLM) and an unsupervised denoising method (Noise2Void[28], N2V). NLM produces an over-smoothed result, and N2V produces a result with patterned noise. By contrast, the HD2Net result denoises the data while preserving most detailed features, demonstrating the superior noise suppression of our method (**Fig. 4c**). Quantitative assessments with PSNR and SSIM are consistent with these visual analyses.

We subsequently applied HD2Net to thicker, multicellular samples. We applied HD2Net to *C. elegans* embryos expressing a GFP-membrane marker labeling head neurons and gut cells imaged with dual-view light-sheet microscopy (diSPIM)[29],[30] (**Fig. 5a, b**). The raw data displays a high degree of noise that obscures membrane boundaries. Compared to raw data, dim features are well resolved in the HD2Net prediction, which enhances resolution and contrast. We found similar improvements when applying HD2Net to *C. elegans* embryos expressing pan-membrane and pan-nuclear labels (**Fig. 5c, d**). In the raw data, many closely spaced membranes and nuclei blur together. HD2Net predictions resolved these targets, further demonstrating the applicability of HD2Net on additional biological samples.

**Fig. 5.**
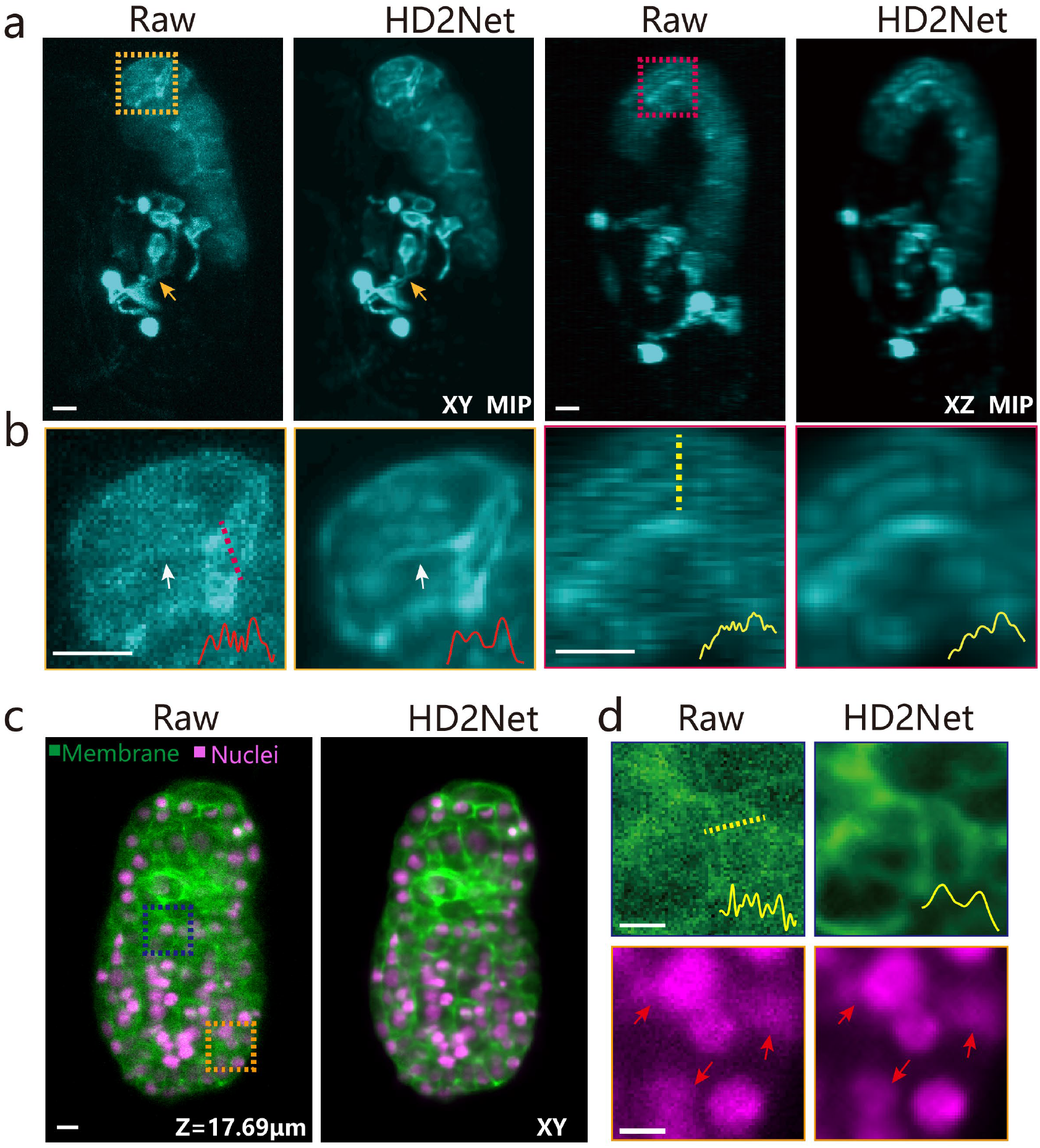
Simultaneous denoising and deaberration improves image quality on different biological samples. **a)** Lateral and axial maximum intensity projection (MIP) images of C. elegans embryos expressing membrane-localized GFP under control of the ttx3-3b promoter, imaged with dual-view light-sheet microscopy (diSPIM), comparing raw recordings and the HD2Net model prediction. Yellow arrows highlight features for visual comparisons. **b)** Higher magnification views of yellow and red rectangular regions in **a**. The insets represent the line profile along the red and yellow lines. White arrows highlight features for visual comparisons. **c)** C. elegans embryos expressing GFP-labeled membrane marker (green) and mCherry-labeled nuclear marker (magenta) were imaged with diSPIM, comparing raw recording (left) and the HD2Net model prediction (right). **d)** Higher magnification views of membranes in the dark blue rectangular region (top) and nuclei in the orange rectangular region (bottom) in **c**. The inset in the top row represents the line profile along the yellow dashed line. The arrows in the bottom row indicate the clear edges of the nuclei recovered by HD2Net. Scalebars: 3 µm in **a** and **b**, 3.5 µm in **c** and **d**.

## 4. Discussion

In this study, we present HD2Net, a powerful computational framework designed to enhance fluorescence microscopy images by simultaneously addressing the challenges posed by optical aberrations and noise. Our findings demonstrate that HD2Net significantly improves image quality compared to existing methods, particularly under low signal-to-noise ratio (SNR) conditions, where traditional approaches struggle. The ability of HD2Net to operate without the need for expensive hardware further enhances its usability for researchers, making high-quality imaging accessible in a wider range of settings[31],[32].

We attribute the improved performance of HD2Net vs. DeAbe or previous denoising methods to two main reasons. First, the integration of a noise estimation module within HD2Net allows for effective noise characterization tailored to the sample, which is crucial when imaging complex biological samples. We directly incorporate this information into the training process, improving subsequent predictions when compared to DeAbe alone. Second, instead of using a single end-to-end restoration structure, we reshape the network architecture by dividing the restoration task to two modules, with each module targeting denoising and deaberration respectively. This design incorporates low-noise images for intermediate supervision, providing more supervisory information that facilitates HD2Net to learn prior knowledge of noise effectively, reducing the difficulty for the following deaberration, and thereby enhancing its overall performance.

While our results are promising, there are still areas for future improvement as well as caveats to the overall method. First, like any deep learning method, HD2Net is fundamentally limited in its ability to reconstruct information that is absent from the raw data, as it can only generate predictions based on the available data. The performance of HD2Net could be further improved by leveraging larger and more diverse datasets for training, which would help the model generalize better to a broader range of imaging conditions. Second, although HD2Net estimates the noise level from the data, its aberration estimation uses a random mixture of relatively low-order aberrations. There is potential for performance improvement through more precise estimation and addition of aberrations. Finally, investigating the application of HD2Net in speed and light dose critical scenarios, such as time-lapse live-cell imaging, could provide insights into the dynamics of biological processes with minimal photodamage [33],[34].

## Code and Data Availability

The source code for HD2Net is available at https://github.com/eguomin/HD2Net/. The code of DeAbe is available at https://github.com/eguomin/DeAbePlus/. Representative data to test the code are publicly available at https://doi.org/10.5281/zenodo.14538488; other datasets are available from the corresponding author upon request due to their large file size.

## Author Contributions

Conceived project: M.G., H.L., and H.S. Developed and implemented HD2Net framework: X.H., Y.L., and M.G. Wrote software: X.H., Y.L. Designed and performed experiments: X.H., M.G., C.M.H. Performed data analysis: X.H. Wrote paper: X.H., Y.L., and M.G. with input from all authors. Supervised research: M.G., H.L., and H.S.

## Acknowledgements

This research was supported by the National Natural Science Foundation of China (62427807, 62475232, 6240030418), the Talent Program of Zhejiang Province (2021R51004), the Postdoctoral Fellowship Program of CPSF (GZB20230626), and the China Postdoctoral Science Foundation (2023M733040). This work was also supported by the Howard Hughes Medical Institute (HHMI). We thank Leanna Eisenman for the PtK2 cell sample preparation; Teng-Leong Chew, the Advanced Imaging Center, and the Light Microscopy facility at HHMI Janelia Research Campus for supporting experiments with the AO-LLSM system. This article is subject to HHMI’s Open Access to Publications policy. HHMI laboratory heads have previously granted a non-exclusive CC BY 4.0 license to the public and a sub-licensable license to HHMI in their research articles. Pursuant to those licenses, the author-accepted manuscript of this article can be made freely available under a CC BY 4.0 license immediately upon publication.

## Reference

[1] J. W. Lichtman and J.-A. Conchello, “Fluorescence microscopy,” Nature methods 2, 910–919 (2005).

[2] Stelzer, “Contrast, resolution, pixelation, dynamic range and signal-to-noise ratio: Fundamental limits to resolution in fluorescence light microscopy,” Journal of Microscopy 189, 15–24 (1998).

[3] A. J. Wright and S. P. Poland, “Adaptive optics for aberration correction in optical microscopy,” in Handbook of Photonics for Biomedical Engineering (Springer Netherlands, 2017), pp. 585–612.

[4] W. Wallace, L. H. Schaefer, and J. R. Swedlow, “A workingperson’s guide to deconvolution in light microscopy,” Biotechniques 31, 1076–1097 (2001).

[5] N. Ji, “Adaptive optical fluorescence microscopy,” Nature methods 14, 374–380 (2017).

[6] K. M. Hampson, R. Turcotte, D. T. Miller, K. Kurokawa, J. R. Males, N. Ji, and M. J. Booth, “Adaptive optics for high-resolution imaging,” Nature Reviews Methods Primers 1, 68 (2021).

[7] D. Gabor, “Microscopy by reconstructed wave-fronts,” Proceedings of the Royal Society of London. Series A. Mathematical and Physical Sciences 197, 454–487 (1949).

[8] M. Guo, Y. Wu, C. M. Hobson, et al., “Deep learning-based aberration compensation improves contrast and resolution in fluorescence microscopy,” Nat. Commun. 16, 313 (2025).

[9] Shroff, H., Testa, I., Jug, F. et al. Live-cell imaging powered by computation. Nat Rev Mol Cell Biol 25, 443–463 (2024).

[10] C. A. Casacio, L. S. Madsen, A. Terrasson, M. Waleed, K. Barnscheidt, B. Hage, M. A. Taylor, and W. P. Bowen, “Quantum-enhanced nonlinear microscopy,” Nature 594, 201–206 (2021).

[11] L. Hu, S. Hu, W. Gong, and K. Si, “Image enhancement for fluorescence microscopy based on deep learning with prior knowledge of aberration,” Opt. Lett. 46, 2055– 2058 (2021).

[12] K. Wang, D. E. Milkie, A. Saxena, P. Engerer, T. Misgeld, M. E. Bronner, J. Mumm, and E. Betzig, “Rapid adaptive optical recovery of optimal resolution over large volumes,” nature methods 11, 625–628 (2014).

[13] V. Ntziachristos, “Going deeper than microscopy: The optical imaging frontier in biology,” Nature methods 7, 603–614 (2010).

[14] L. N. Thibos, R. A. Applegate, J. T. Schwiegerling, and R. Webb, “Standards for reporting the optical aberrations of eyes,” (2002).

[15] N. Dey, L. Blanc-Feraud, C. Zimmer, et al., “Richardson–lucy algorithm with total variation regularization for 3d confocal microscope deconvolution,” Microsc. research technique 69, 260–266 (2006).

[16] M. F. Freeman and J. W. Tukey, “Transformations related to the angular and the square root,” The annals of mathematical statistics 607–611 (1950).

[17] M. Makitalo and A. Foi, “Optimal inversion of the anscombe transformation in low-count poisson image denoising,” IEEE transactions on Image Processing 20, 99–109 (2010).

[18] F. J. Anscombe, “The transformation of poisson, binomial and negative-binomial data,” Biometrika 35, 246–254 (1948).

[19] X. Liu, M. Tanaka, and M. Okutomi, “Noise level estimation using weak textured patches of a single noisy image,” in 2012 19th IEEE International Conference on Image Processing (IEEE, 2012), pp. 665–668.

[20] S. Wold, K. Esbensen, and P. Geladi, “Principal component analysis,” Chemometrics and intelligent laboratory systems 2, 37–52 (1987).

[21] J. Chen, H. Sasaki, H. Lai, Y. Su, J. Liu, Y. Wu, A. Zhovmer, C. A. Combs, I. Rey-Suarez, H.-Y. Chang, and others, “Three-dimensional residual channel attention networks denoise and sharpen fluorescence microscopy image volumes,” Nature methods 18, 678–687 (2021).

[22] Y. Zhang, K. Li, K. Li, L. Wang, B. Zhong, and Y. Fu, “Image super-resolution using very deep residual channel attention networks,” in Proceedings of the European Conference on Computer Vision (ECCV) (2018), pp. 286–301.

[23] P. Charbonnier, L. Blanc-Feraud, G. Aubert, and M. Barlaud, “Two deterministic half-quadratic regularization algorithms for computed imaging,” in Proceedings of 1st International Conference on Image Processing (IEEE, 1994), Vol. 2, pp. 168–172.

[24] Y. Wu, X. Han, Y. Su, M. Glidewell, J. S. Daniels, J. Liu, T. Sengupta, I. Rey-Suarez, R. Fischer, A. Patel, and others, “Multiview confocal super-resolution microscopy,” Nature 600, 279–284 (2021).

[25] B.-C. Chen, W. R. Legant, K. Wang, L. Shao, D. E. Milkie, M. W. Davidson, C. Janetopoulos, X. S. Wu, J. A. Hammer Iii, Z. Liu, and others, “Lattice light-sheet microscopy: Imaging molecules to embryos at high spatiotemporal resolution,” Science 346, 1257998 (2014).

[26] T.-L. Liu, S. Upadhyayula, D. E. Milkie, V. Singh, K. Wang, I. A. Swinburne, K. R. Mosaliganti, Z. M. Collins, T. W. Hiscock, J. Shea, and others, “Observing the cell in its native state: Imaging subcellular dynamics in multicellular organisms,” Science 360, eaaq1392 (2018).

[27] A. Buades, B. Coll, and J.-M. Morel, “Non-local means denoising,” Image Processing On Line 1, 208–212 (2011).

[28] A. Krull, T.-O. Buchholz, and F. Jug, “Noise2void-learning denoising from single noisy images,” in Proceedings of the IEEE/CVF Conference on Computer Vision and Pattern Recognition (2019), pp. 2129–2137.

[29] Wu, Y., Wawrzusin, P., Senseney, J. et al. Spatially isotropic four-dimensional imaging with dual-view plane illumination microscopy. Nat Biotechnol 31, 1032–1038 (2013). 10.1038/nbt.2713.

[30] Y. Wu, A. Kumar, C. Smith, E. Ardiel, P. Chandris, R. Christensen, I. Rey-Suarez, M. Guo, H. D. Vishwasrao, J. Chen, and others, “Reflective imaging improves spatiotemporal resolution and collection efficiency in light sheet microscopy,” Nature communications 8, 1452 (2017).

[31] Y. Li, Y. Su, M. Guo, X. Han, J. Liu, H. D. Vishwasrao, X. Li, R. Christensen, T. Sengupta, M. W. Moyle, and others, “Incorporating the image formation process into deep learning improves network performance,” Nature Methods 19, 1427–1437 (2022).

[32] M. Guo, Y. Li, Y. Su, T. Lambert, D. D. Nogare, M. W. Moyle, L. H. Duncan, R. Ikegami, A. Santella, I. Rey-Suarez, and others, “Rapid image deconvolution and multiview fusion for optical microscopy,” Nature biotechnology 38, 1337–1346 (2020).

[33] A. Mishin and K. Lukyanov, “Live-cell super-resolution fluorescence microscopy,” Biochemistry (Moscow) 84, 19–31 (2019).

[34] A. G. Godin, B. Lounis, and L. Cognet, “Super-resolution microscopy approaches for live cell imaging,” Biophysical journal 107, 1777–1784 (2014).

